# A Proximity Sensor-Based Multipurpose Object Recognition Test System for Rodent Behavior Research

**DOI:** 10.1101/2022.11.29.518044

**Authors:** Octavio G.G. Vazquez, Mehmet A. Demirhan, Mackenzie Bannister, Qian-Quan Sun

**Affiliations:** Wyoming Sensory Biology Center, University of Wyoming; Department of Zoology and Physiology, University of Wyoming; Department of Mechanical Engineering, University of Wyoming

## Abstract

We report an automated, low-cost, and open-source methodology for rodent object recognition tasks (ORT), which are widely used behavior paradigms. The system was designed with an effort of behavior standardization and improvement. When it combined with easily accessible 3-D manufacturing technology, this proximity capacitive sensor based method provides an opportunity to explore new, and high throughput hardware for NORT. The system’s software allows behavior data to be instantaneously monitored and its output can be plotted promptly immediately upon the completion of the behavior test, thus saving labor and time. The performance of this package is cross-verified with a widely used video-based automated software scoring method and examined its sensitivity to chemogenetic behavior manipulations. This integrated package is an affordable and accurate tool for translational recognition memory research and other type of research with frequent use of NORT paradigms.

**Visual Abstract:** 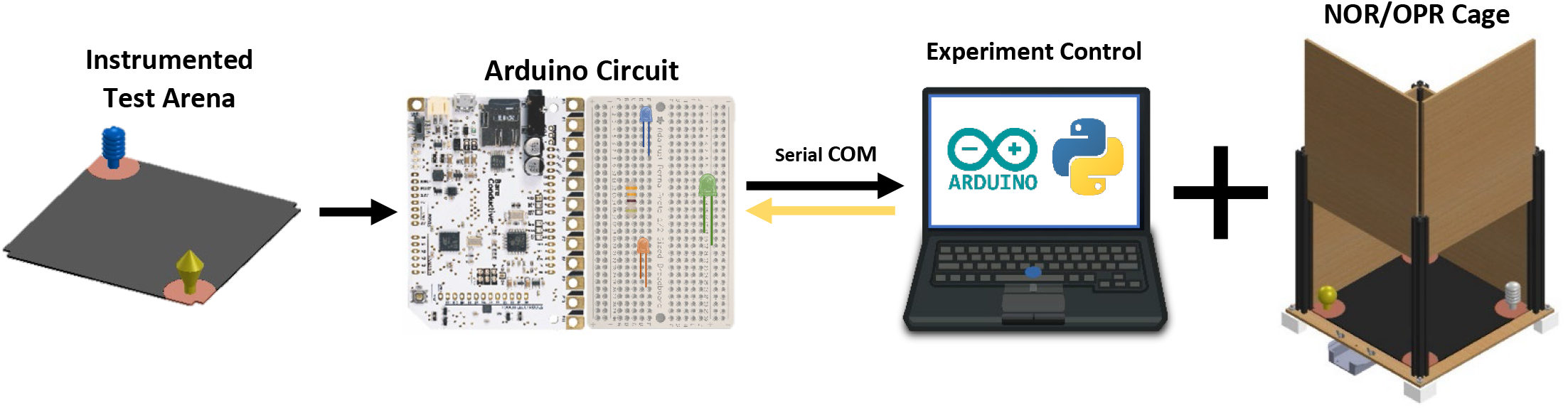

## Introduction

Rodents tend to spend more time with a novel object than with a familiar object (Berlyne, 1954, 1966). This tendency to novel object has been shown to be dependent on sensory dependent cognitive memories and also correlated to the level of anxiety in rodents (Bevins and Besheer, 2006; Ennaceur and Delacour, 1988). A behavior test paradigm, known as novel object recognition test (NORT) has been developed to measure animal’s recognition memory (Antunes and Biala, 2012; Bevins and Besheer, 2006). The NORT has rapidly and increasingly become a widely used behavior paradigm because it does not require extensive animal training, exposure to aversive stimuli or food or water deprivation, and tests can be executed in one session (Antunes and Biala, 2012; Bevins and Besheer, 2006). Using the key word ‘NORT’ to perform a Pubmed search yield nearly 15,000 published papers, of which many use NORT results as a proxy for recognition memory and translational research (Antunes and Biala, 2012; Bevins and Besheer, 2006; Ennaceur and Delacour, 1988; Grayson et al., 2014; Heyser and Chemero, 2012)

With variant parameters such as the object size, material, and exact location, as well as different scoring approaches, object recognition tasks (ORTs) are performed technically differently across labs (Antunes and Biala, 2012). The main ORTs experiments are the NORT and Object Place Recognition Test (OPRT), and the relevant data acquired from them include, but are not limited to, the test subject cumulative time and entries investigating the different objects (Lueptow, 2017). Previously, while time consuming, trained observers would live score these experiments manually using a stopwatch (Clarke et al., 2010; Williams et al., 2007). However, OTRs are currently mostly assessed with camera recordings and visual software technology. Different methods are used in video analysis of behavior experiments. In order of simplicity – the first one is to score behavior on the videos entirely manually (Denninger et al., 2018). The second one is to use a behavior sequence event-log software, to assist detailed video and audio manual scoring (Noldus et al., 2000). The third one, and most widely used, is to use a video analysis software with some degree of automation. There is robust open-source software for automated behavior video analysis. For instance, different mouse behaviors at its home-cage have been successfully identified(Jhuang et al., 2010), as well as different social interactions(de Chaumont et al., 2012). However, most of open-source programs are usually tailored to a specific-type of experiment or application, and the source-code is often too time-consuming to properly learn and adapt, leaving many neuroscientists to prefer user-friendly, commercially made softwares.

For many rodent behavioral experiments’ mazes and ORTs, the most popular visual analysis software is Noldus’ EthosVision, which uses nose-point tracking, and is as accurate as a trained observer(Benice and Raber, 2008). Noldus equipment, however, comes with a significant price tag, and require videos to be taken under perfect visual conditions (e.g. lighting and contrasting color between animal and background). EthosVision will require lengthy manual tracking corrections (Benice and Raber, 2008). For the afore-mentioned constraints, capacitive proximity sensors for behavioral experiments data acquisition have been researched for more than a decade(Badelt and Blaisdell, 2008), and in recent years, it was offered as a reliable, simple, and affordable alternative. Several low-cost microcontroller peripherals offer multiple touch/capacitive sensors for various applications. A widely commercially available one is the MPR121 chip, a sensor controller that features 12 independent smart electrodes, which is offered with online code repositories that allow for different charge sensitivity detection, and other configurations. The first report of the MPR121 used for a behavioral task was on tampered beam test, where it showed a 91.6% accuracy (Ardesch et al., 2017). Later, under the name of CapTouch, this same instrumentation was proposed to perform ORTs, using the investigation object as the sensor electrode itself, while taking advantage of widely available 3D printing equipment to standardize shapes and sizes of the respective objects(Spry et al., 2021).

Searching for a way to easily implement, test, and troubleshoot the MPR121 capacitive sensors, we designed a cost effective and simple data acquisition system, and protocol, which uses only Arduino computers, and open software, which we name and present as Capacitive Plate Sensor System, CAPSS. Moreover, we provide source hardware materials for an individual Object Recognition Cage Test Arena, which optimizes material usage. The CAPSS is exclusively manufactured using 3D printers, and laser cutting, which is available for most university and research labs. This cage is compact, easy to clean, handle, and can be repurposed to perform other experiments or to accommodate test subjects. We provide validation of the sensitivity of CAPSS for NORT, using a variety of manipulations including chemogenetic manipulations and genetic strains known to affect NORT. We provide parallel comparisons to automated video scoring approaches.

## Materials and Methods

### Components and Construction

We designed a cage arena to easily tinker hardware and software improvements. The cage requirements were that the walls are tall enough to prevent the mouse from escaping, a design that allowed for easy cleaning after experiments, and an elevated floor for careful wiring. After its assembly, miscellaneous lab tasks allowed different versions of this cage to be developed to fulfill specific purposes as shown in Figure 2.D. We share this hardware to highlight the potential of laser cutting and 3D printing to effectively resolve small accommodation issues behavioral neuroscientists face often. The cage total volume is 15’ by 15” and 18” tall, and the arena’s 12’’ by 12’’ and 15’’ tall. We share all its construction files and more in *#Repository*.

**Table 1.** Component List for Two Multipurpose Cage Arenas with Correspondent CAPSS System.

| Component | Quantity | Supplier | ID | Total Price |
| --- | --- | --- | --- | --- |
| 48" x 48" x 1/4" Medium Density Fiberboard Sheet | 1 | McMaster | 2726N67 | \$ 33.72 |
| 12" x 24" x 1/8" Black Cast Acrylic Sheet | 1 | McMaster | 8505K745 | \$ 15.40 |
| 12" x 12" 0.01" UV-Resistant Acrylic Film | 1 | McMaster | 4076N11 | \$ 6.31 |
| Pack of 25. 1/4"-20 Thread Size, 1" Long Stainless Steel Thread-Locking Socket Head Screw | 1 | McMaster | 93705A542 | \$ 10.01 |
| Pack of 100. 1/4" - 20. Medium-Strength Steel Hex Nuts—Grade 5 | 1 | McMaster | 95462A029 | \$ 7.81 |
| Pack for 100. Stainless Steel Washer for 1/4" Screw Size, 0.281" ID, 0.625" OD | 1 | McMaster | 92141A029 | \$ 3.47 |
| Pack for 100. Stainless Steel Button Head Hex Drive Screw 8-32 Thread Size, 1/2" Long | 1 | McMaster | 92949A194 | \$ 8.51 |
| Pack of 25. Stainless Steel Wing Nut 8-32 Thread Size | 1 | McMaster | 92001A291 | \$ 6.84 |
| XE25L15 - 25 mm Square Construction Rail, 15" Long, 1/4"-20 Taps | 8 | ThorLabs | XE25L15 | \$ 171.28 |
| Copper Foil Tape with Conductive Adhesive - 25mm x 15 meter roll | 2 | Adafruit | 1127 | \$ 39.90 |
| Adafruit Perma-Proto Half-sized Breadboard PCB - Single | 2 | Adafruit | 1609 | \$ 9.00 |
| Solder Wire - 60/40 Rosin Core - 0.5mm/0.02" diameter - 50 grams | 1 | Adafruit | 1886 | \$ 5.95 |
| Short Headers Kit for Feather - 12-pin + 16-pin Female Headers | 4 | Adafruit | 2940 | \$ 6.00 |
| 2.54mm/0.1" Pitch Terminal Block - 2-pin | 4 | Adafruit | 2138 | \$ 3.80 |
| Diffused 10mm LED Pack - 5 LEDs each in 5 Colors - 25 Pack | 1 | Adafruit | 4204 | \$ 7.95 |
| Blue Wire Spool | 1 | Adafruit | 2989 | \$ 2.95 |
| Red Wire Spool | 1 | Adafruit | 288 | \$ 2.95 |
| Bare Conductive Arduino Touch Board | 2 | eduporium | 5013 | \$ 115.90 |
| USB micro-B Cable - 6 Foot | 2 | Sparkfun | 10215 | \$ 11.00 |
| Prusament PLA Gentleman's Grey 1kg | 1 | Prusa | PRM-PLA-GMG- | \$ 24.99 |
| | | | TOTAL | \$ 493.74 |

**Figure 1.**
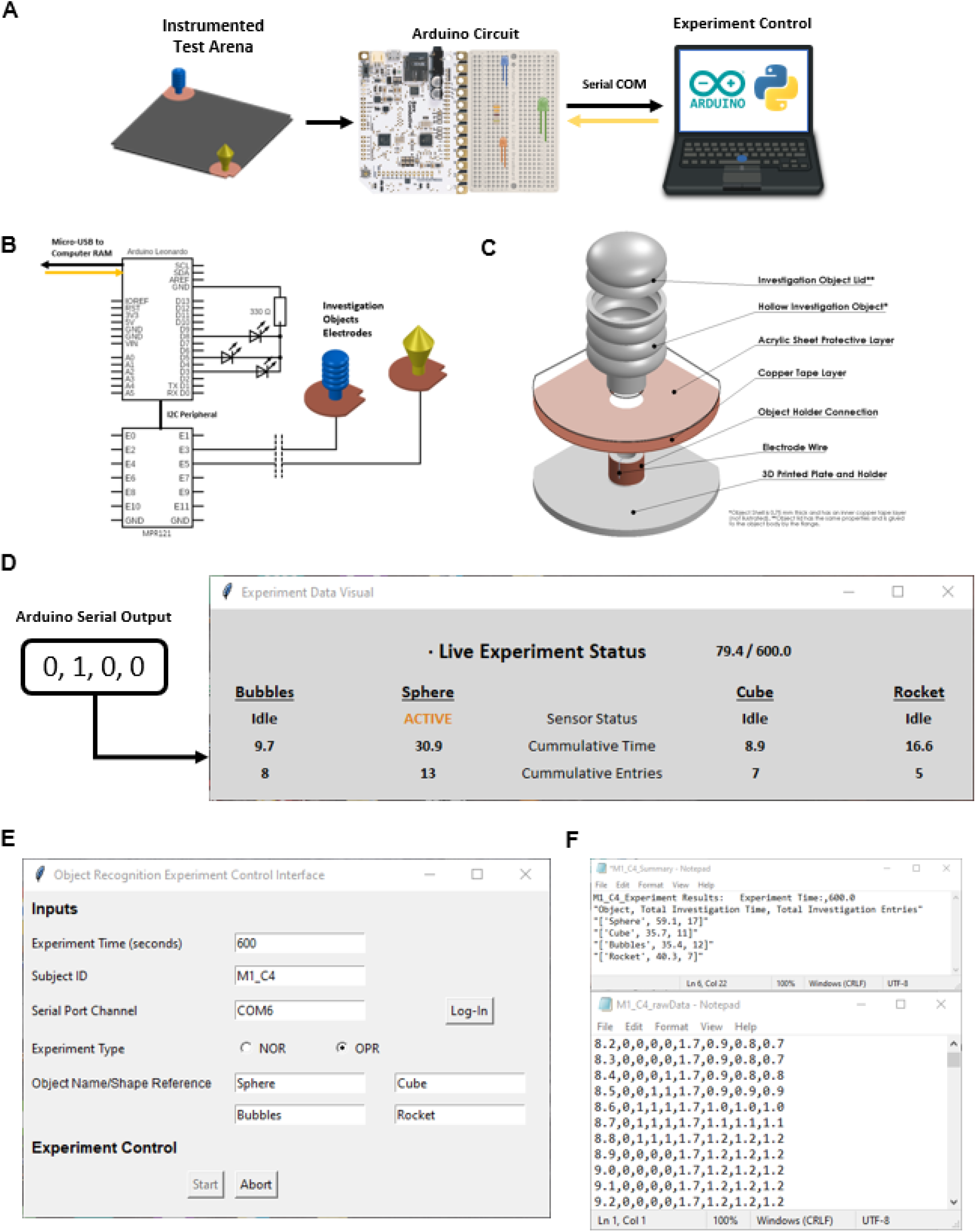
CAPSS overview and features **A.** Data acquisition overview. **B.** Arduino circuit diagram for NORT. **C.** Sensor plate and object holder for ORTs. **D.** Live data visual of OPRT experiment. **E.** User inputs and experiment control GUI. **F.** CAPSS data output after finalized experiment.

**Figure 2.**
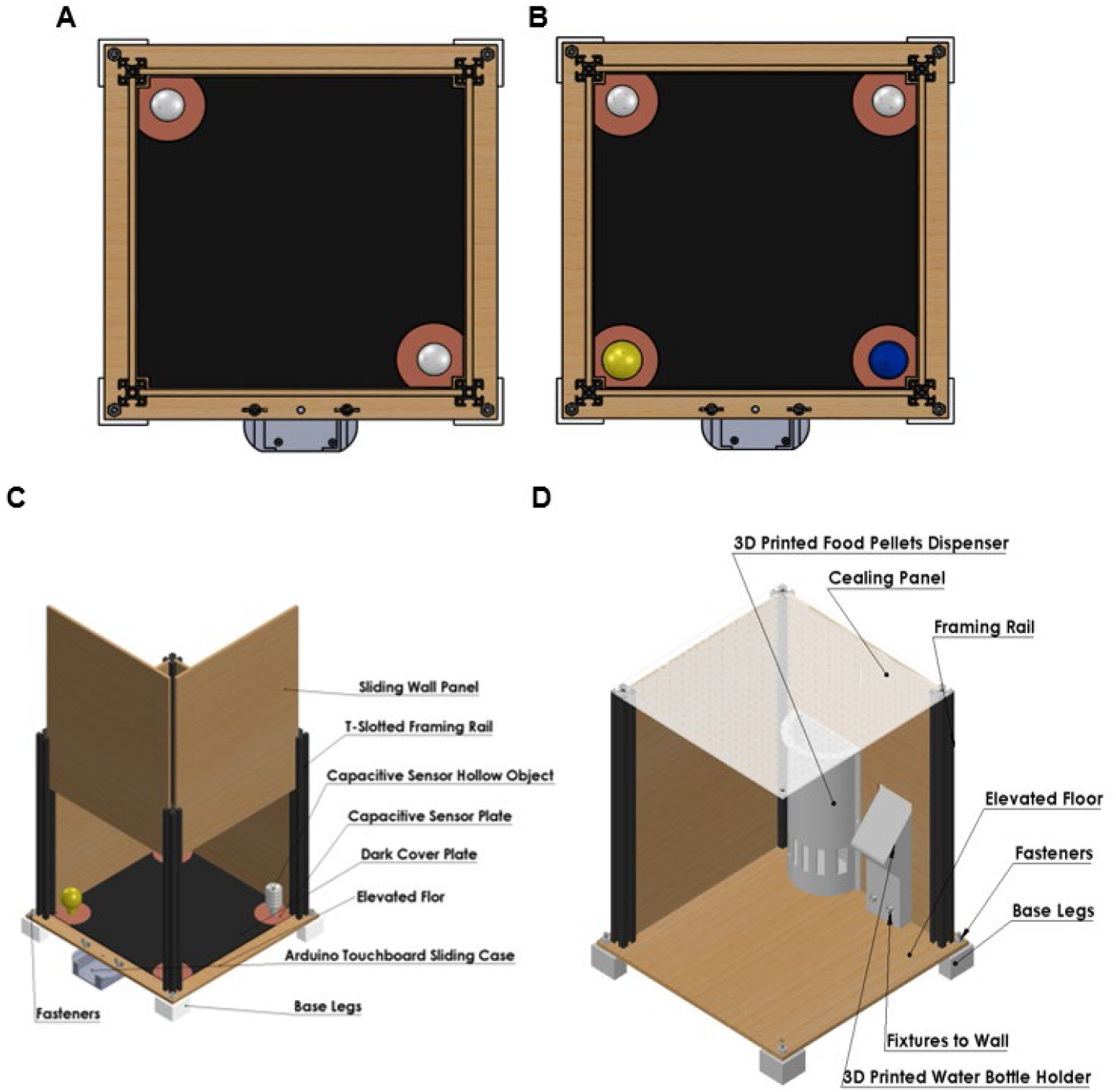
Multipurpose Cage Arena **A.** Novel Object Recognition Set-up **B.** Object Place Recognition Set-Up **C.** Cage parts and features **D.** Open cage view of safe setup for mouse with Microdrive implant.

### Testing Procedure

Our main goal was to compare the performance of CAPSS (sensor plates and objects) to CapTouch (only sensor objects), and then compare both to automated video score using a widely used software (Noldus’ EthosVision). With simultaneous video recording, we performed six, NOR-testing-phase experiments (no familiarization or habituation), under bright light conditions, with each respective hardware configuration (CAPSS and CapTouch). The video was recorded by a Balser acA1300-60gm camera, with a 1.5 - 12.5mm, 1:1.2 Computar Lens. The NOR investigation objects were hollow and had an inner layer of copper tape, made from different color PLA, had dimensions of less than 1.5” x 1.5” x 2.5”, and both were determined as climbable (they are shown in Figure 1.A and 1.B). The behavior tests were conducted double blindly by two lab members, each performing only one set of tests. First, the video recording was started, then the mouse test subject was placed on the NOR arena, and immediately the sensor data acquisition system was started to sample for the experiment duration. After each trial, the cage arena was cleaned and sanitized with 70% ethanol; and before a new trial, the Arduino with its peripheral MPR121 was reset to allow sensor charge sensitivity re-calibration. Each NOR experiment had two object sensors, meaning twelve data samples were collected per hardware system. Lastly, to further show sensitivity of CAPSS in measuring short-term memory changes in the rodents, we examined CAPSS NORT in animals treated with chemogenic activation and silencing of the Papez circuits (Rolls, 2019).The protocol for these experiments was the same as the described above.

### Animals

All experiments were conducted according to protocols approved by the Institutional Animal Care and Use Committee (IACUC) of the University of Wyoming. Mice cages were maintained in a room where 22–23 °C on a 12:12 h light-dark cycle. Food and water were available ad libitum. Ank1b mutant mice(Lopez et al., 2017) and a wild-type mice were used and obtained from Jackson Laboratories (Bar Harbor, ME). For the CAPPS/CapTouch/EthosVision comparison experiments, four three-months-old wild-type ANK mice and 2 Ank1b mutant mice were used. To prevent order effect, half of the animals tested first with plates then tested with no plates while the other half tested first with no plates and then with plates. For chemogenetic activation and silencing experiments two months old Thy1-JRGECO1a mice were tested. Activation group (n=6) received pAAV-hSyn-hM3D(Gq)-mCherry (AAV9) injection and silencing group (n=6) received pAAV-hSyn-DIO-hM4D(Gi)-mCherry (AAV8) injection to their Anterior Thalamus region and location of the virus were validated with histology.

### Analysis

We compared the cumulative time spent in each object and its percentages generated by Ethovision and CAPSS, respectively (Fig 3). The effective time samples were obtained from the videos, using Noldus’ EthosVision’s nose-point tracking video analysis in the “any nose or center point in investigation zone” configuration, since we were not discriminating against climbing. And from the sensors, the effective data was summarized from the raw experiment data samples using a MATLAB script. For chemogenetic activation and silencing experiments, only summarized raw CAPSS data was used. All processed data statistics, per system, was computed, plotted, and compared, using Excel.

**Figure 3.**
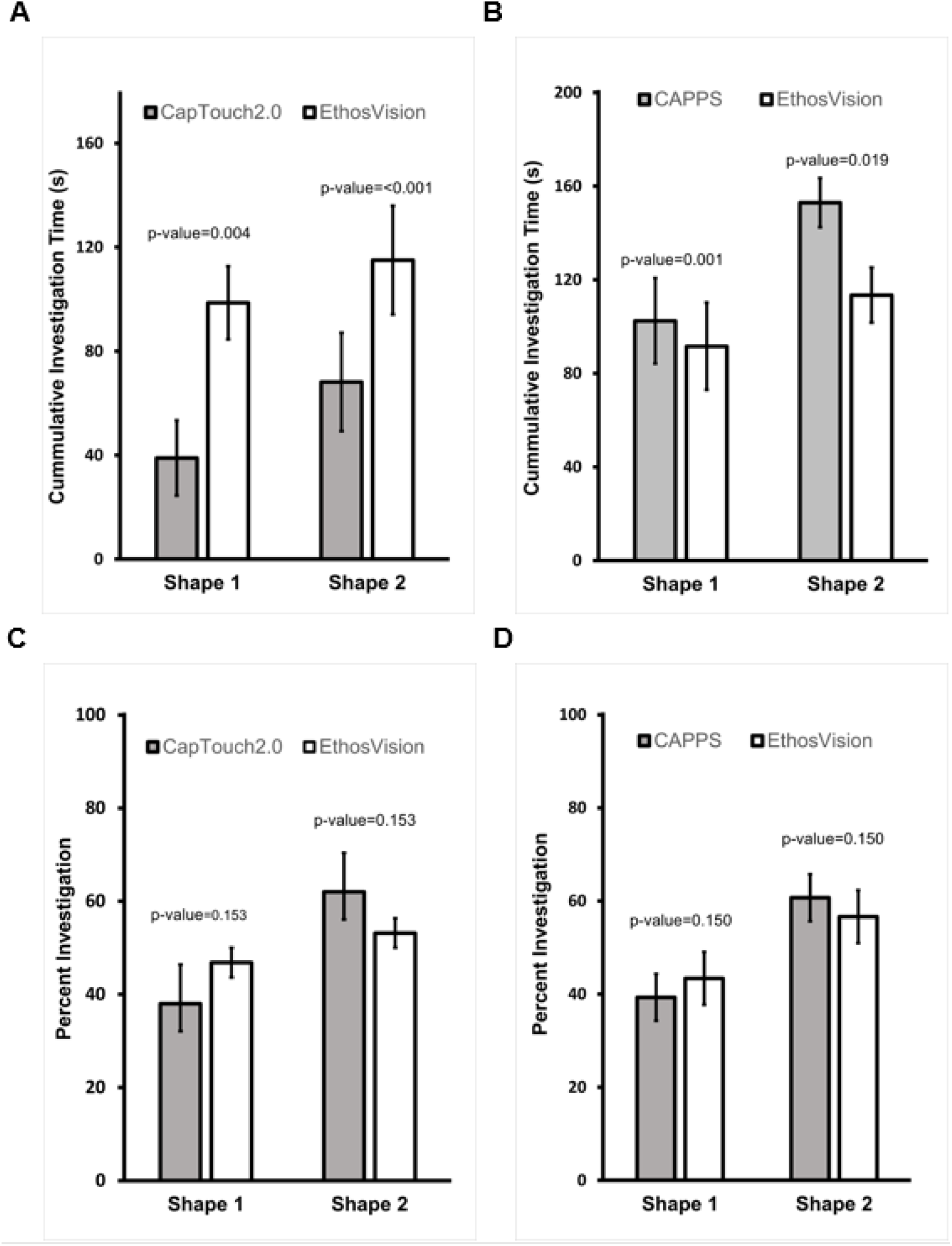
Sensor Hardware Performance Comparison. **A.** CapTouch Average Investigation Time per Object Compared to EthosVision’s. Object 1 had a time score difference of −59.67 seconds, and Object 2, −46.88 seconds. **B.** CAPSS Average Object Investigation Time per Object Compared to EthosVision’s. The difference was +10.90 seconds and +39.49 seconds, for each object, respectively. **C.** CapTouch Average Investigation Percentage per Object Compared to EthosVision’s. The difference was 8.82% **D.** Average Investigation Percentage per Object Compared to EthosVision’s. The difference was 4.08%

## Results

Overview of CAPSS performing a NORT experiment. A custom Arduino with an embedded MPR121, called TouchBoard, is used to instrument a test arena, and sequentially connected to a computer through its micro-USB port (Fig 1A). The microcontroller data acquisition and the experiment are controlled from the computer through a Python open-source graphic user interface (GUI). This GUI is easy to use and allows – experiment log inputs (Fig 1E), live monitoring (Fig 1D), and results and raw data storage (Fig 1F). We also offer another monitoring and data output alternative using only Arduino code in its Integrated Development Environment. We provide all source materials in the GitHub repository (*https://github.com/wyomingneuron/CAPSS*).

The main hardware difference, compared to previous capacitive object sensors, is shown in Figure 1.C. To detect the proximity induced capacitance related current caused by mouse approaching to one of the objects, the MPR121 sensor controller must run at its highest code-configurable sensitivity, and even then, it requires the test subject to get into direct contact with the sensor surface. To detect an investigation event entry, the mouse needs to touch, lean, or climb the sensor object, and this often discriminates against soft entries such as close looking and sniffing. To solve this problem, we added a surrounding electrode plate that allows detection when the mouse is close by the object, without requiring contact with it. CAPSS is then a hollow object with an interior layer of copper tape (like CapTouch, but) connected to a conductive plate.

We observe how CapTouch underscores, while CAPSS over-scored investigation time compared to EthosVision (Figure 3). However, in both configurations, computed investigation percentage (object investigation time*100/total investigation time, Figure 3.C and 3.D), showed results similar to EthosVision results. The output of CAPSS raw data is an experiment sample every 0.1 seconds, consisting of - the current experiment time, sensor statutes, and their current chronometer time value. Plotting these parameters produces the graphs seen in Figure 4. A, B and C, which are used to gain a glance into locomotion patterns such as frequency and duration of investigation events. To evaluate if the CAPSS can detect behavior changes caused by the effects of chemogenetic silencing or activation of the anterior thalamus, a key node of the Papez circuit (Rolls, 2019), known to be critical for anxiety mode and memories, we performed NORT in naïve untreated mice and in mice treated with synthetic ligand Clozapine-N-Oxide (CNO) injection (1 mg/kg) (Armbruster et al., 2007). Results from Figure 4.D demonstrates how data acquired from CAPSS yields statistical significance in NORT when we manipulate anterior thalamic circuit necessary for the NORT task.

**Figure 4.**
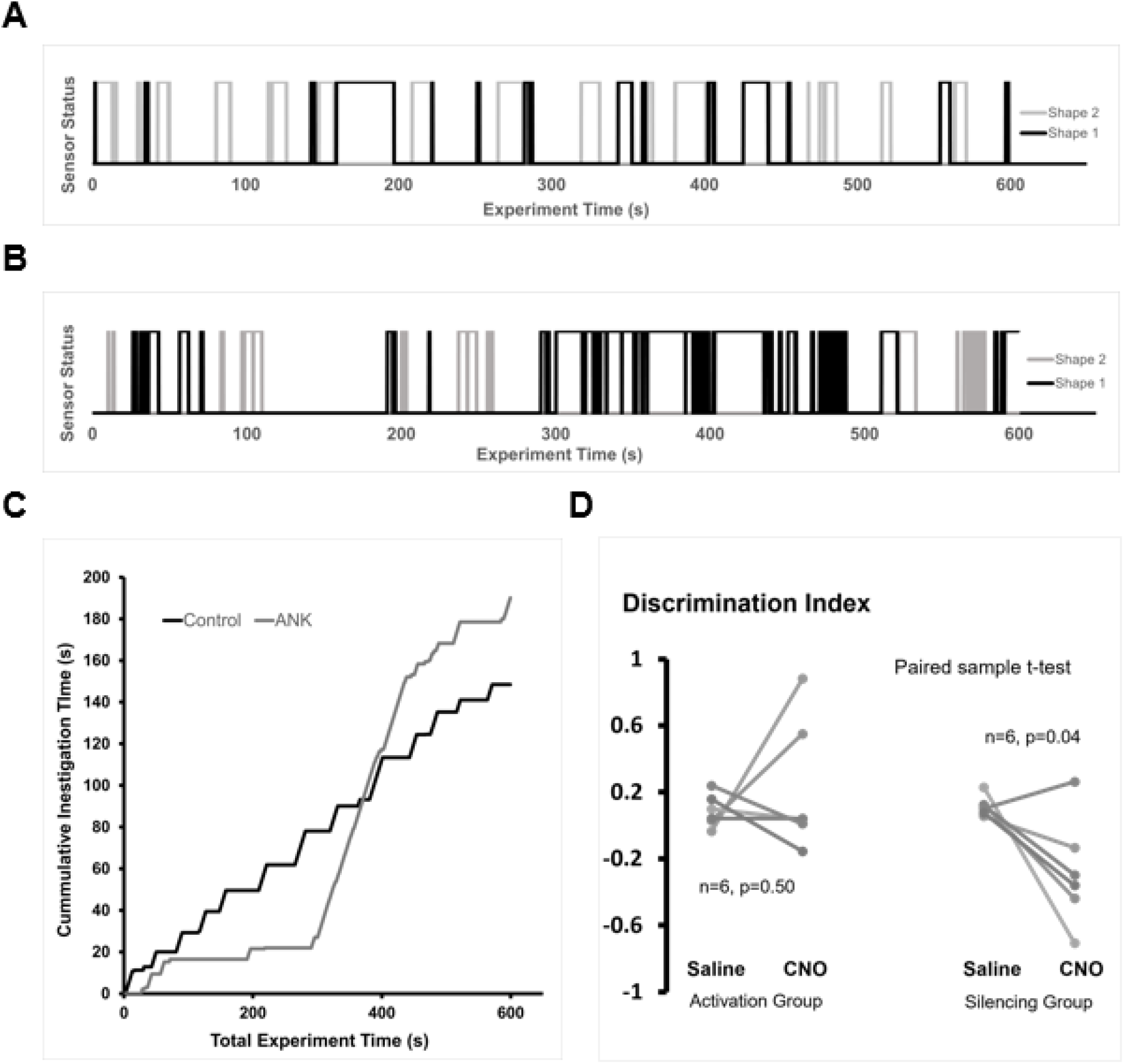
Raw CAPSS NOR Data Plotted and Chemogenetics NOR Experiment Results. **A.** CAPSS NOR Experiment Entries for a Control Mouse. **B.** CAPSS NOR Experiment Entries for ANK Mouse. **C.** CAPSS NOR Cumulative Investigation Time Comparison for Shape 1. **D.** Chemogenetically manipulated animal’s discrimination index scores after intraperitoneal injection of saline and CNO injection.

## Discussion

First, we have presented CAPSS, an open-source, low-cost, and effective approach for data acquisition on object recognition tasks. CAPSS consists of sensor microcontrollers outputting directly into the computer’s serial COM ports, and monitoring and storing the data using open-source software. Second, we have showcased the potential of additive manufacturing, and laser cutting to quickly tackle miscellaneous laboratory equipment set-up challenges. Third, we have re-introduced the design for rodent experiment capacitive sensors, which is in the form of plates that the test subject can step on(Badelt and Blaisdell, 2008). We compared CAPSS against visual analysis for NOR-like experiments, and yielded a less than a 5% object investigation time percentage average differences. With the current design, these sensors do not distinguish nose vs. tail orientation of the test subject. However, from the 12-sample video analysis, we found that only 12% of the total zone entries were tail entries with little investigations, which explains a source of difference in investigation time. Overall, capacitive sensor systems offer various practical advantages over visual analysis, the most significant one being cost-effectiveness, particularly in the time required for analysis. Using the field’s most cited video tracking software, Ethovision, as comparison, without perfect lighting conditions, it will need labor intensive manual scoring for video frames. Even under the best achievable bright light arena conditions, the software missed tracking of the 3-point mouse orientation for several frames, and to correct this error, it added up to about 15 hours manual analysis for the twelve trials. Moreover, if all conditions are optimized and manual tracking correction can be avoided, capacitive sensors are still attractive for its data size. Without accounting all supporting files for the visual analysis, the size of the twelve experiment videos was 5.56GB, while the total size of all sensor data was only 1.53MB. Finally, capacitive sensors are mostly attractive, for their programable TTL pulses, Figure 1.B, that can allow a simultaneous input for in-vivo behavioral maze experiments data acquisition, which is difficult for the visual systems to do.

### System Limitations

It was mentioned that to operate, the capacitive sensors must run at their highest sensitivity. This makes them susceptible to detect noise. When troubleshooting CAPSS (and CapTouch), a recurring issue was far, but active DC wires and human presence, that would trigger the electrode falsely. Further, the electrode wires cannot be shielded since it makes them lose sensitivity enough to stop detecting the test subject, meaning that for the MPR121, in room conditions, the noise magnitude that gets grounded, is about the same or more of the electrostatic charge of a mouse. The solution for this was first, to place the arenas in a faraday cage; and second, to mount the Arduino computer with its MPR121 sensors, underneath the arena cage, in a 3D printed, PLA enclosure box.

### Future Directions

First, as mentioned, NORTs need standardization. Both arena size and object characteristics should be standardized, and minimum parametric requirements in the different experiment phases should be defined. For investigation objects, as proposed in Fry et al.’s study (2021), different shape 3D printable object models could quickly create a common object repository for ORTs; and we offer our optimized cage, or at least its arena surface dimensions, 12’’ by 12’’, as a standard for mice ORTs. Second and last, regarding object recognition, the capacitive sensors’ accuracy could be improved by splitting the investigation object and surrounding plates into many electrodes. This would allow more data acquisition to compute the specific location of the mouse around, behind, or on top of the object, and then discriminate incorrect investigation entries or climbing.

